# Complete Genome Sequence of the Polysaccharide-Degrading Rumen Bacterium *Pseudobutyrivibrio xylanivorans* MA3014

**DOI:** 10.1101/2020.07.08.194233

**Authors:** Nikola Palevich, Paul H. Maclean, William J. Kelly, Sinead C. Leahy, Jasna Rakonjac, Graeme T. Attwood

## Abstract

Ruminants are essential for maintaining the global population and managing greenhouse gas emissions. In the rumen, bacterial species belonging to the genera rumen *Butyrivibrio* and *Pseudobutyrivibrio* constitute the core bacterial rumen microbiome and are important degraders of plant-derived complex polysaccharides. *Pseudobutyrivibrio xylanivorans* MA3014 was selected for genome sequencing in order to examine its ability to breakdown and utilize plant polysaccharides. The complete genome sequence of MA3014 is 3.58 Mb, consists of three replicons (a chromosome, chromid and plasmid), has an overall G+C content of 39.6% and encodes 3,265 putative protein-coding genes (PCGs). Comparative pan-genomics of all cultivated and currently available *P. xylanivorans* genomes has revealed highly open genomes and a strong correlation of orthologous genes within this species of rumen bacteria. MA3014 is metabolically versatile and capable of utilizing a range of simple mono-or oligosaccharides to complex plant polysaccharides such as pectins, mannans, starch and hemicelluloses for growth, with lactate, butyrate and formate as the principal fermentation end-products. The genes encoding these metabolic pathways have been identified and MA3014 is predicted to encode an extensive repertoire of Carbohydrate-Active enZYmes (CAZymes) with 80 Glycoside Hydrolases (GHs), 28 Carbohydrate Esterases (CEs) and 51 Glycosyl Transferases (GTs), that suggest its role as an initiator of primary solubilization of plant matter in the rumen.

## Introduction

*Butyrivibrio* and *Pseudobutyrivibrio* represent the most commonly isolated butyrate-producing anaerobic rumen bacteria (Henderson, et al. 2015), and are among the small number of rumen genera capable of utilizing the complex plant structural polysaccharide xylan (Bryant and Small 1956; Hungate 1966). *Pseudobutyrivibrio* [family Lachnospiraceae, order Clostridiales] are anaerobic, monotrichous, butyrate-producing, curved rods and have been isolated from the gastrointestinal tracts of various ruminants, monogastric animals and humans (Kopecný, et al. 2003; Willems and Collins 2009). The *Butyrivibrio* and *Pseudobutyrivibrio* genera originally consisted of only one species, *Butyrivibrio fibrisolvens* (Bryant and Small 1956). In addition to phenotypic characterisations (Hazlewood, et al. 1986; Shane, et al. 1969), studies have utilized DNA-DNA hybridization (Mannarelli 1988; Mannarelli, et al. 1990), 16*S* rRNA gene sequencing (Forster, et al. 1996; Willems, et al. 1996) and 16*S* rRNA-based hybridization probes (Forster, et al. 1997), to differentiate these organisms. To accommodate the observed diversity amongst the newly discovered bacterial strains, a new genus, *Pseudobutyrivibrio*, was described in which only *P. ruminis* and *P. xylanivorans* species are currently recognized (Kopecný, et al. 2003; Van Gylswyk, et al. 1996). *P. xylanivorans* are common anaerobic rumen bacteria found in domestic and wild ruminants and the type strain is Mz 5^T^ (DSM 14809) (Henderson, et al. 2015; Kopecný, et al. 2003). *P. xylanivorans* Mz 5^T^ is non-proteolytic but is able to utilize xylan or hemicellulose and various oligo- and monosaccharides as substrates for growth (Zorec, et al. 2000). Gaining an insight into the role of these microbial primary plant polysaccharide fermenters is important for understanding rumen function. Here we present the complete genome sequence of *P. xylanivorans* MA3014, a strain isolated from a New Zealand pasture-grazed dairy cow (Seshadri, et al. 2018), and describe its comparison with other representative *P. xylanivorans* genomes.

## Materials and Methods

### Growth Conditions and Fermentation End Product Analysis

*P. xylanivorans* MA3014 was isolated from the rumen contents of fistulated Friesian dairy cattle and sequenced (Noel 2013; Seshadri, et al. 2018). MA3014 was grown in RM02 medium (Kenters, et al. 2011) with 10 mM glucose and 0.1% yeast extract but without rumen fluid and culture purity was confirmed by Gram stain. The morphological features of MA3014 cells were determined by both scanning (SEM) and transmission (TEM) electron microscopy of cells grown on RM02 medium alone or with the addition of neutral detergent fraction (NDF) of plant material as previously described (Palevich, et al. 2017; Palevich, et al. 2018).

Growth on soluble substrates was assessed as an increase in culture density OD_600nm_ compared to cultures without carbon source added (all tested at 0.5% w/v final concentration), whereas total VFA production was used as an indicator of substrate utilization and growth for insoluble polymers (Supplementary Table S3). VFA production was determined from triplicate broth cultures grown overnight in RM02 medium with cellobiose as substrate and analysed for formate, acetate, propionate, n-butyrate, iso-valerate and lactate on a HP 6890 series GC (Hewlett Packard) with 2-ethylbutyric acid (Sigma-Aldrich, St. Louis, MO, USA) as the internal standard. To derivatize formic, lactic and succinic acids, samples were mixed with HCl ACS reagent (Sigma-Aldrich, St. Louis, MO, USA) and diethyl ether, with the addition of N-methyl-N-t-butyldimethylsilyltri-fluoroacetamide (MTBSTFA) (Sigma-Aldrich, St. Louis, MO, USA) (Richardson, et al. 1989).

### Preparation of Genomic DNA for Whole-Genome Sequencing

Genomic DNA was extracted from freshly grown cells by a modification of the standard cell lysis method previously described (Palevich, et al. 2018; Seshadri, et al. 2018), followed by phenol-chloroform extraction, and purification using the Qiagen Genomic-Tip 500 Maxi kit (Qiagen, Hilden, Germany). Specificity of genomic DNA was verified by automated Sanger sequencing of the 16*S* rRNA gene following PCR amplification from genomic DNA. Total DNA amounts were determined using a NanoDrop^®^ ND-1000 (Thermo Scientific Inc.) and a Qubit Fluorometer dsDNA BR Kit (Invitrogen, USA), in accordance with the manufacturer’s instructions. Genomic DNA integrity was verified by agarose gel electrophoresis and using a 2000 BioAnalyzer (Agilent, USA).

### Genome Sequencing, Assembly and Comparison

*Pseudobutyrivibrio xylanivorans* MA3014 was selected for genome sequencing as a NZ strain and only representative member of *P. xylanivorans* from the Hungate1000 collection ((Seshadri, et al. 2018): Supplementary Table S1). The complete genome sequence of MA3014 was determined by pyrosequencing 3 kb mate paired-end sequence libraries using the 454 GS FLX platform with Titanium chemistry (Macrogen, Korea). Pyrosequencing reads provided 55× coverage of the genome and were assembled using the Newbler assembler (version 2.7, Roche 454 Life Sciences, USA) which resulted in 116 contigs across 13 scaffolds. Gap closure was managed using the Staden package (Staden, et al. 1999) and gaps were closed using additional Sanger sequencing by standard and inverse PCR techniques. In addition, MA3014 genomic DNA was sequenced using shotgun sequencing of 2 kb paired-end sequence libraries using the Illumina MiSeq platform (Macrogen, Korea) which provided 677-fold sequencing coverage. A *de novo* assembly was performed using the assemblers Velvet version 3.0 (Zerbino and Birney 2008), and EDENA version 3.120926 (Hernandez, et al. 2008). The resulting sequences were combined with the Newbler assembly using the Staden package and Geneious, version 8.1 (Kearse, et al. 2012). Genome assembly was confirmed by pulsed-field gel electrophoresis (Palevich 2011; Palevich N, et al. 2019b) and genome annotation was performed as described previously (Kelly, et al. 2010). Genome comparisons of orthologous gene clusters within *Pseudobutyrivibrio* genomes were performed using OrthoVenn version 2 (Wang, et al. 2015).

## Results and Discussion

### Genome Assembly, Properties and Annotation

To sequence the genome of *P. xylanivorans* MA3014, short-read 454 GS FLX Titanium and Illumina technologies based on 9.9 million paired-end (PE) reads was applied (Supplementary Tables S1 and S2). The genome of *P. xylanivorans* MA3014 consists of three replicons (Palevich 2011, 2016); a single chromosome (3,412,851 bp, %G+C 39.7), a chromid or secondary chromosome (PxyII, 88,942 bp, %G+C 36.9) and a plasmid (pNP95, 82,698 bp, %G+C 37.4) (Figure 1A). The total size of the closed genome is 3,584,491 bp with an overall %G+C content of 39.6%. The MA3014 assembly with high coverage of 677× was achieved using insert sizes that ranged between 238□bp (Illumina MiSeq) and 2.5kb (454 GS-FLX Titanium). In total, 2.6 Gb of trimmed and filtered sequence data was retained for the reported assembly (Supplementary Table S2). The overall genome assembly statistics of MA3014 are similar to the Mz 5^T^ (DSM 14809) and NCFB 2399 (DSM 10317) ((Kopečný, et al. 2003): Supplementary Table S4).

*Ab initio* gene prediction resulted in a total of 3,365 genes annotated in MA3014, of which 3,265 (97.03%) were PCGs (Supplementary Table S4). Among these, a putative function was assigned to 2,364 (70.25%), while 601 PCGs were annotated as hypothetical proteins or proteins of unknown function. In total, 840 (24.96%) genes have clear homology to proteins in the KEGG database, 2,506 (74.47%) and 2,593 (77.06%) of annotated genes have well-defined PFAM and InterPro protein domains, respectively. In contrast, 153 (4.55%) of annotated genes have identified signal peptide protein domain hits and are predicted have extracellular functions. The MA3014 chromosome encodes 3,098 PCGs while the PxyII and pNP95 encode 96 and 71 genes, respectively. Overall, the coding region comprises 89.77% of the genome, typical of rumen bacterial genomes (Seshadri, et al. 2018; Palevich, et al. 2019b). However, in order to elucidate the actual genetic divergence within the rumen *Butyrivibrio* and *Pseudobutyrivibrio*, future efforts should focus on the generation of complete genomes, starting with the Hungate1000 collection.

**Figure 1.**
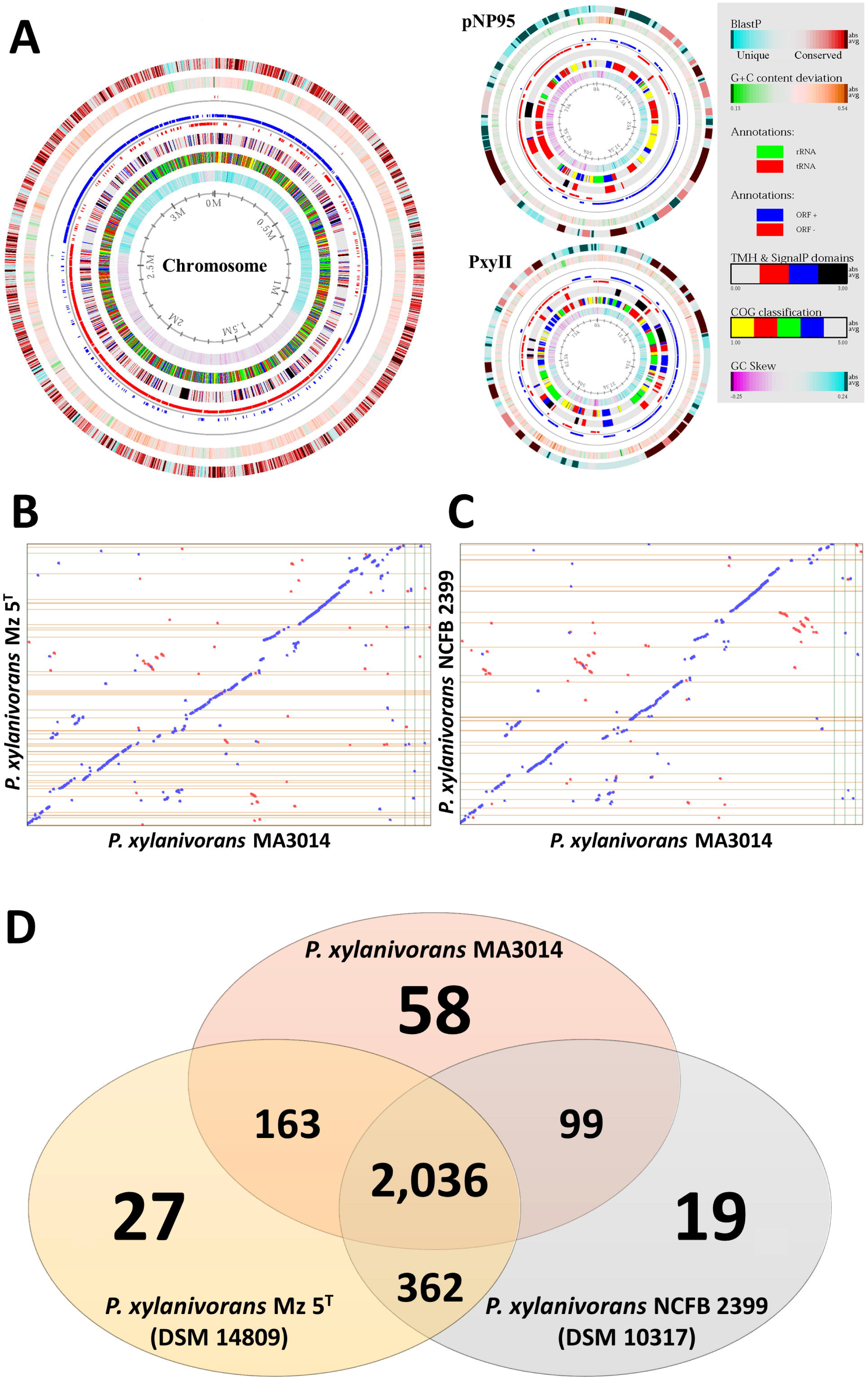
(A) Genome atlas for *P. xylanivorans* MA3014. The figure represents a circular view of the four replicons that constitute the *P. xylanivorans* MA3014 genome. The key at the right describes the concentric circles within each replicon in the outermost to innermost direction. Circle 1 (innermost circle) indicates GC-skew. Circle 2 shows COG classifications of predicted and annotated open reading fames (ORFs) grouped into five major categories: information storage and processing (yellow); cellular processes and signalling (red); metabolism (green); poorly characterized (blue); uncharacterized or no COG assignment (uncoloured). Circle 3 shows transmembrane helices (TMH) and SignalP domains grouped into four categories: both absent (uncoloured); TMH (red); SignalP (blue); both present (black). Circle 4 indicates ORF orientation in sense (ORF+, blue) or antisense (ORF-, red) directions. Circle 5 shows tRNA (green) and rRNA (red) ribosomal machinery. Circle 6 shows G+C content deviation from the average in either green (low GC spike) or orange (high GC spike). Circle 7 shows BLAST similarities of unique proteins (blue) and highly conserved features (red) relative to sequences in the nonredundant (nr) database. (B-C) Genome synteny analysis. Alignment of the *P. xylanivorans* MA3014 genome against the draft genomes of *P. xylanivorans* Mz 5^T^ (B) and *P. xylanivorans* NCFB 2399 (C). Whenever the two sequences agree, a colored line or dot is plotted. Units displayed in base-pairs. Color codes: blue, forward sequence, red, reverse sequence. (D) Venn diagram showing the distribution of shared gene families among the *P. xylanivorans* genomes. All *P. xylanivorans* scaffolds with at least a single one-to-one ortholog shared among the genomes were compared.

### Genome Comparison

A comparison of the *P. xylanivorans* MA3014 genome with the draft genomes of *P. xylanivorans* Mz 5^T^ (DSM 14809) and NCFB 2399 (DSM 10317) (Kopečný, et al. 2003) is shown in Supplementary Table S4. MA3014 is the largest *P. xylanivorans* genome to date, where it is 163,567 bp and 370,547 bp larger, also contains 187 and 375 more PCGs than Mz 5^T^ and NCFB 2399, respectively. A novel feature of MA3014 and other well-characterized *Butyrivibrio* genomes is the presence of chromids or secondary chromosomes (Kelly, et al. 2010; Palevich, et al. 2019a). Chromids are replicons with %G+C content similar to that of their main chromosome, but have plasmid-type maintenance and replication systems, are usually smaller than the chromosome (but larger than plasmids) and contain genes essential for growth along with several core genus-specific genes (Harrison, et al. 2010). The PxyII replicon has been designated as a chromid of MA3014 as it possesses all of these characteristics and contains genes encoding enzymes that have a role in carbohydrate metabolism and transport. Since the PxyII chromid of MA3014 is 2,834 bp smaller than the Bhu II chromid of MB2003, it is now the smallest chromid reported for bacteria. Although several plasmid replication genes have been identified in the Mz 5^T^ but not in NCFB 2399 draft genomes, the presence of extrachromosomal elements requires experimental validation.

Comparison of MA3014, Mz 5^T^ and NCFB 2399 genomes based on COG category (Supplementary Table S5) and synteny analysis (Figure 1B-C), show that these *Pseudobutyrivibrio* strains are genetically similar. Despite the differences in genome sizes of MA3014 and Mz 5^T^, the basic metabolism of these two rumen bacteria are comparable. Comparative pan-genomics of these rumen bacterial strains revealed highly open genomes and a strong correlation of orthologous genes among these species (Figure 1D). Most of the predicted MA3014 genes were found to have homologs (BLASTP e-value cut-off 10^−5^) in the other two strains (2,356; 73%), with the *P. xylanivorans* represented by 768 orthologous clusters and 1,996 single-copy genes. In total, 2,036 core genes were found to be orthologous among the three *P. xylanivorans* genomes compared, with only 58 genes found to be unique to MA3014 (Figure 1D). Genomic comparisons with other species within the genera *Butyrivibrio* and *Pseudobutyrivibrio* have revealed strong collinearities (Palevich, et al. 2019b), that will facilitate our understanding of genome evolution of rumen bacteria.

### Polysaccharide Degradation

The Carbohydrate-Active enZYmes database was used to identify glycoside hydrolases (GHs), glycosyl transferases (GTs), polysaccharide lyases (PLs), carbohydrate esterases (CEs) and carbohydrate-binding protein module (CBM) families within the MA3014 genome. Overall, CAZyme profile of MA3014 is similar to other *Pseudobutyrivibrio* that are in general not as extensive as those of *Butyrivibrio* (Palevich 2016; Palevich, et al. 2019b). Analysis of the functional domains of enzymes involved in the breakdown or synthesis of complex carbohydrates, has revealed the polysaccharide-degrading potential of this rumen bacterium (Supplementary Table S6). Approximately 2% of the MA3014 genome (69 CDSs) is predicted to encode either 22 secreted (21 GHs and one CE) and 47 intracellular (41 GHs, 4 CEs and two GTs) proteins dedicated to polysaccharide degradation. The enzymatic profiles of MA3014 and Mz 5^T^ are almost identical, as both possess the same genes encoding predicted secreted and intracellular CAZymes in their genomes. Out of the 22 genes predicted to encode secreted polysaccharide degrading enzymes, only β-glucosidase *bgl3K* (FXF36_15770) is encoded by the MA3014 chromid (PxyII). The majority (40) of MA3014 genes encoding intracellular proteins involved in polysaccharide breakdown (excluding GTs), had corresponding homologues in Mz 5^T^. The most abundant Pfam domains included GH families (GH3, GH13 and GH43) and CE1, most of which did not contain signal sequences and predicted to be located intracellularly. Similarly, CAZymes with predicted roles in xylan (GH8, GH51, GH115), dextrin and starch (GH13 and GH77) degradation families were also predicted to be mostly intracellular.

Growth experiments showed MB2003 to be a metabolically versatile bacterium able to grow on a wide variety of monosaccharides and disaccharides (Supplementary Table S3). However, unlike Mz 5^T^ (Kopecný, et al. 2003), MA3014 was unable to utilize the insoluble substrate pectin for growth. This difference is due to Mz 5^T^ possession of 4 pectate lyases (1 PL1 and 3 PL3) predicted to be involved in pectin degradation and utilization, of which MA3014 has none. The ability of MA3014 to breakdown starch and xylan is predicted to be based on four large (>1,000 aa) cell-associated proteins shown to be significantly up-regulated in related *B. hungatei* MB2003 and *B. proteoclasticus* B316^T^ cells grown on xylan (Kelly, et al. 2010; Palevich, et al. 2019a). These are: α-amylase *amy13E* (FXF36_11320), arabinogalactan endo-1,4-β-galactosidase *agn53A* (FXF36_02635), xylosidase/arabinofuranosidase *xsa43D* (FXF36_08285), endo-1,4-β-xylanase xyn10A (FXF36_14365). These proteins contain multiple cell wall binding repeat domains (CW-binding domain, Pfam01473) at their C-termini that are predicted to anchor the protein to the peptidoglycan cell membrane (Dunne, et al. 2011). In addition, the secreted α-amylase *amy13E* (FXF36_11320) contains a CBM26 (Pfam16738) domain with predicted starch-binding functions (Gilbert, et al. 2013; McCartney, et al. 2004).

Electron microscopy of MA3014 cells grown in liquid media supplemented with plant material has revealed the copious production of exopolysaccharides (EPS) (Figure 2A-B). EPS is a characteristic of *Butyrivibrio* strains and is composed of the neutral sugars rhamnose, fucose, mannose, galactose and glucose (Stack 1988), made available by recycling the plant polysaccharides breakdown products. Our findings also show the presence of cytoplasmic inclusions (Figure 2C), similar to those seen in B316^T^ and other *Butyrivibrio* strains containing glycogen-like material (Hespell, et al. 1993). The MA3014 genome encodes a complete repertoire of genes for glycogen synthesis and degradation, suggesting that a variety of complex oligosaccharides resulting from extracellular hydrolysis are metabolized within the cell and that glycogen has a role in the storage of excess carbohydrate.

**Figure 2.**
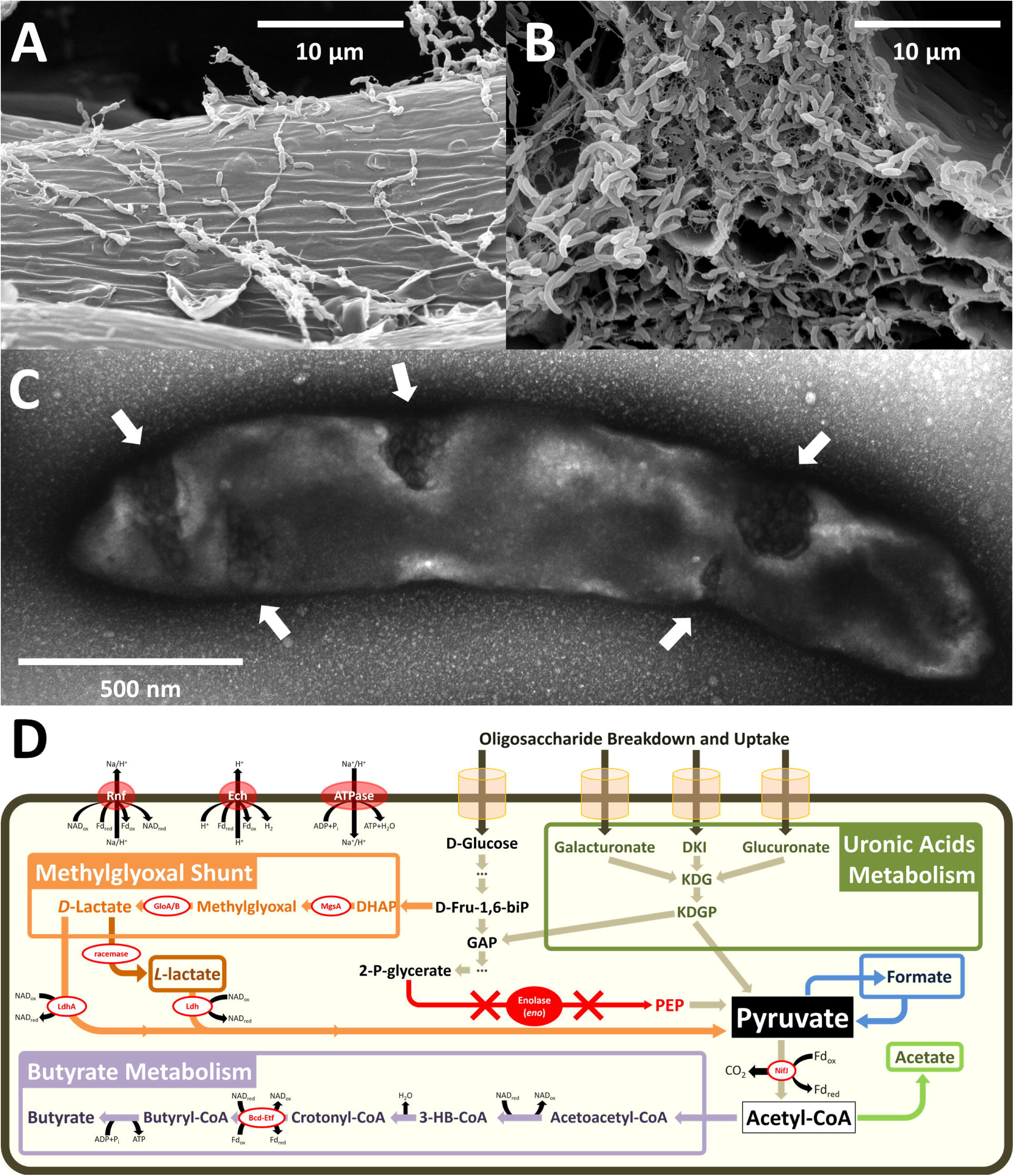
(A-C) Electron micrographs of *P. xylanivorans* MA3014. (A-B) Scanning EMs of MA3014 cells adherence to the surface (A) and exposed end (B) of NDF plant material, at 5,000 × magnification. (C) Transmission EM of negatively stained MA3014 cells grown in liquid medium at 10,000 × magnification. Arrows indicate the presence of glycogen inclusions. (D) Fermentation pathways in rumen *Pseudobutyrivibrio* and *Butyrivibrio*. Abbreviations: Bcd-Etf, butyryl-CoA dehydrogenase/electron transferring flavoprotein; Ech, *E. coli* hydrogenase-3-type hydrogenase; Fd, ferredoxin; Fd_ox_, oxidized Fd; Fd_red_, reduced Fd; Glo, glyoxalase; MsgA, methylglyoxal synthase; NAD, nicotinamide adenine dinucleotide; NAD_ox_, oxidized NAD; NAD_red_, reduced NAD; NifJ, nitrogen fixation J; Rnf, *Rhodobacter* nitrogen fixation; ATPase = F_0_F_1_-ATPsynthase.

### Enolase Loss and Metabolic Flexibility

*Pseudobutyrivibrio* and *Butyrivibrio* are key members of the degradative microbiota found in a highly carbohydrate-rich environment and appear to have evolved beyond using glycolysis as the central pathway. The pathways for butyrate production for these rumen bacteria presume the possession of a complete Embden-Meyerhof-Parnas (EMP) glycolytic pathway. The lack of an enolase (*eno*, EC4.2.1.11), that converts 2-phospho-D-glycerate to phosphoenolpyruvate in the second to last step of the EMP pathway, is extremely unusual. As part of the Hungate1000 project in which the genomes of 410 rumen microbes were sequenced (Seshadri, et al. 2018), we discovered that many specialized polysaccharide fermenters lacked an enolase gene. Of all 21 *Pseudobutyrivibrio* genomes sequenced, only *P. xylanivorans* MA3014 and *P. ruminis* AD2017 reported to date lack a detectable enolase. *Eno-P. xylanivorans* MA3014 and several *Butyrivibrio* strains were recently confirmed using PCR screens with *eno*-specific primers (Kelly, et al. 2010; Palevich, et al. 2018). Given the observed polysaccharide-degrading abilities and lactate production, the Methylglyoxal Shunt (MS) and uronic acid metabolic pathways (Figure 2D), have been suggested as alternatives to the EMP pathway (Cooper 1984). Previous work has also reported similar findings in other *Butyrivibrio* strains (Kelly, et al. 2010; Palevich, et al. 2019a). In this pathway the dihydroxyacetone phosphate is transformed to pyruvate via methylglyoxal and D-lactate dehydrogenase encoded by *ldhA*. The MA3014 genome possesses methylglyoxal synthase, *mgsA* (FXF36_12340), glyoxalases *gloA/B* (FXF36_00730, FXF36_01130 and FXF36_09530) and both D- and L-lactate dehydrogenases *ldh* (FXF36_04170 and FXF36_11135) genes. In addition, MA3014 has the same set of genes as the previously reported and well-characterized *B. hungatei* MB2003 and *B. proteoclasticus* B316^T^ for the production of butyrate, formate, acetate and lactate (Kelly, et al. 2010; Palevich, et al. 2019a; Palevich, et al. 2017).

In some butyrate-forming anaerobes, crotonyl-CoA reduction is linked to electron transport phosphorylation (ETP) via flavin-based electron bifurcating *ech* and *rnf* complexes which act as transmembrane ion pumps (Buckel and Thauer 2013; Herrmann, et al. 2008; Li, et al. 2008; Welte, et al. 2010). A recent analysis of the Hungate1000 dataset (Hackmann and Firkins 2015; Seshadri, et al. 2018; Palevich, et al. 2019b), found that *Pseudobutyrivibrio* and *Butyrivibrio* genomes encode both Ech and Rnf homologues proposed to act in concert with NifJ and Bcd-Etf to form an electrochemical potential and drive ATP synthesis (Gutekunst, et al. 2014; Tremblay, et al. 2013). This allows these rumen bacteria to generate approximately 4.5 ATP/glucose in total, one the highest yields for anaerobic fermentation of glucose (Buckel and Thauer 2013). Given the importance of *eno, Pseudobutyrivibrio* and *Butyrivibrio* may be displaying an example of environment-specific evolution by gene loss that warrants further investigation into the alternative pathways that permit ATP generation. The genome sequence of *P. xylanivorans* MA3014 presented here is consistent with the genome architecture of other sequenced *Pseudobutyrivibrio* strains and is a valuable resource for future studies regarding bacterial-driven plant-fibre degradation in ruminants.

## Supplementary Material

Supplementary data are available at *Genome Biology and Evolution* online.

### Data deposition

The complete genome sequence of *Pseudobutyrivibrio xylanivorans* MA3014 and its annotations are deposited in Genbank under accession numbers CP043028, CP043029 and CP043030.

## Acknowledgements

The *P. xylanivorans* MA3014 genome sequencing project was funded by the New Zealand Ministry of Business, Innovation and Employment New Economy Research Fund programme: Accessing the uncultured rumen microbiome, contract number C10×0803. Electron microscopy was conducted with the assistance of the Manawatū Microscopy and Imaging Centre at Massey University, Palmerston North, New Zealand. We thank Sarah Lewis for assistance with the fermentation end product analysis, Abdul Baten and Ron Ronimus for critical review of the manuscript.

